# Ultrastructural Dynamics of Dopaminergic Presynaptic Release Sites revealed by Cryo-correlative Light and Electron Microscopy

**DOI:** 10.1101/2024.04.15.589543

**Authors:** Matthew Domenic Lycas, Dustin R. Morado, Ulrik Gether, John A.G. Briggs, Simon Erlendsson

## Abstract

Dopaminergic neurons are fundamental in governing motivation, movement, and many aspects of cognition. The targeted modulation of dopaminergic signaling serves as a cornerstone in developing therapeutic interventions for conditions such as Parkinson’s disease, schizophrenia, and addiction. Despite the pivotal role of dopaminergic neurons, the ultrastructure and associated dynamics of dopaminergic synapses remain poorly understood. Here, we develop and utilize a cryo-correlative light and electron microscopy process chain to investigate the micro- to nanoscale architecture and organelle content of dopaminergic presynaptic release sites. Using cryo electron tomography, we identify several protein complexes crucial to dopaminergic function and we utilize subtomogram averaging to resolve *in situ* assemblies of the TRiC/CCT chaperone and vacuolar-type ATPase. Lastly, we find that pharmacological treatments using either dopamine or the dopamine D2 receptor antagonist, haloperidol, bidirectionally modulate vesicular content, mitochondrial size and calcium phosphate deposition. These findings contribute to our general understanding of the composition and ultrastructural dynamics of dopaminergic presynaptic release sites and provide a methodological platform for further studies of the structure and cell biology of dopaminergic neurons and their responses.

## Introduction

Midbrain dopamine (DA) neurons encode movement, motivation, and reward prediction error through DA release from axons targeting the basal ganglia, the limbic system, and prefrontal cortex. Communication from a given DA neuron is facilitated by the release of neurotransmitters from presynaptic release sites which are often defined as swellings along the axon (varicosities) containing the machinery for neurotransmitter release^1^. Despite their central role in the treatment of various neurological and psychiatric disorders, such as Parkinson’s disease (PD), schizophrenia and drug addiction^2^, very little is known about the molecular architecture and ultrastructure of DA presynaptic release sites. Key components have been identified through proteomic analyses by mass spectroscopy, which offers temporal and compositional detail^3,4^. Spatial organization has been obtained using light microscopy and serial sectioning electron microscopy that have provided insights into the DA neuronal connectome and (re)organization of specific protein components following pharmacological induced changes^5–11^. Together, these methods have provided a general understanding of the content and structure of DA neurons, however, they are decoupled, spatially limited in resolution and depend on chemical fixation which influences neuronal integrity^12^. For hippocampal glutamatergic, GABAergic, and dorsal root ganglia (DRG) neurons, cryo-fixation and subsequent electron microscopy and tomography (cryoEM and cryoET, respectively) have offered high resolution ultrastructural information^13–15^. However, such studies are lacking for primary DA neurons. These require support from glial cells to grow, making it challenging to create a suitable environment in which DA neuronal cultures develop.

In this study, we used a co-culture setup combined with cryo-correlative light and electron microscopy (cryoCLEM) allowing for a multiscale characterization of primary cultured DA neurons. At the micrometer to higher nanometer scale, we gained understanding of which synaptic organelles are represented in DA presynaptic release sites as well as their frequency and distribution. At the lower nanometer scale, we obtained three-dimensional information on organelle connectivity and identified larger molecular structures including voltage gated K^+^/Na^+^/Ca^2+^ channels, the T-complex protein Ring Complex/Chaperonin Containing TCP-1 (TRiC/CCT) and the Vacuolar-type ATPase (V-ATPase)^16,17^. Furthermore, from the tomograms, we resolved low resolution *in situ* structures of both TRiC/CCT and the V-ATPase. Finally, we applied pharmacological stimulation using DA or haloperidol, which resulted in changes in mitochondria size and calcium phosphate content, as well as the number of synaptic vesicles. Our findings represent unprecedented insights into the composition and plasticity of DA presynaptic release sites, shedding light on the structural aspects and molecular components involved in DA release.

## Results

For imaging of primary DA neurons by cryoEM we developed a “Cell-Grid-Cell” preparation. We placed an EM grid on a support layer of cortical glial cells and seeded midbrain neurons on top of the grid. In this way, the grid is sandwiched between the two cell layers. We were only able to promote growth of midbrain neurons in the presence of the supporting cortical glial cell layer (Figure 1A). Upon removal of the grid from the culture, only few glial cells originating from the support cell layer remained attached to the grid, thereby leaving predominantly the midbrain neurons on top of the grid for subsequent cryo fixation and imaging (Figure S1). To identify the DA neurons in the mixed midbrain neuronal population, which also contains non-DA neurons and glia cells, we used positive fluorescence labeling. DA neurons were labelled by transducing the neurons with two adeno-associated viruses (AAVs). The first virus contains the genetic information for the floxed fluorescent protein, tdTomato, and the second virus for cre-recombinase under the control of the tyrosine hydroxylase (TH) promoter^7^. TH catalyzes the conversion of tyrosine into L-DOPA which is a precursor for DA and serves as a marker of DA presynaptic release sites. Following transduction, we were able to observe tdTomato fluorescence both live and following cryo-preservation (Figure S2). From the cryo fixated samples, fluorescence and electron microscopy atlases were collected and correlated. This approach allowed for identification of TH-positive DA neuronal extensions and presynaptic release sites in the mixed midbrain neuronal cultures (Figure 1B-C).

**Figure 1.**
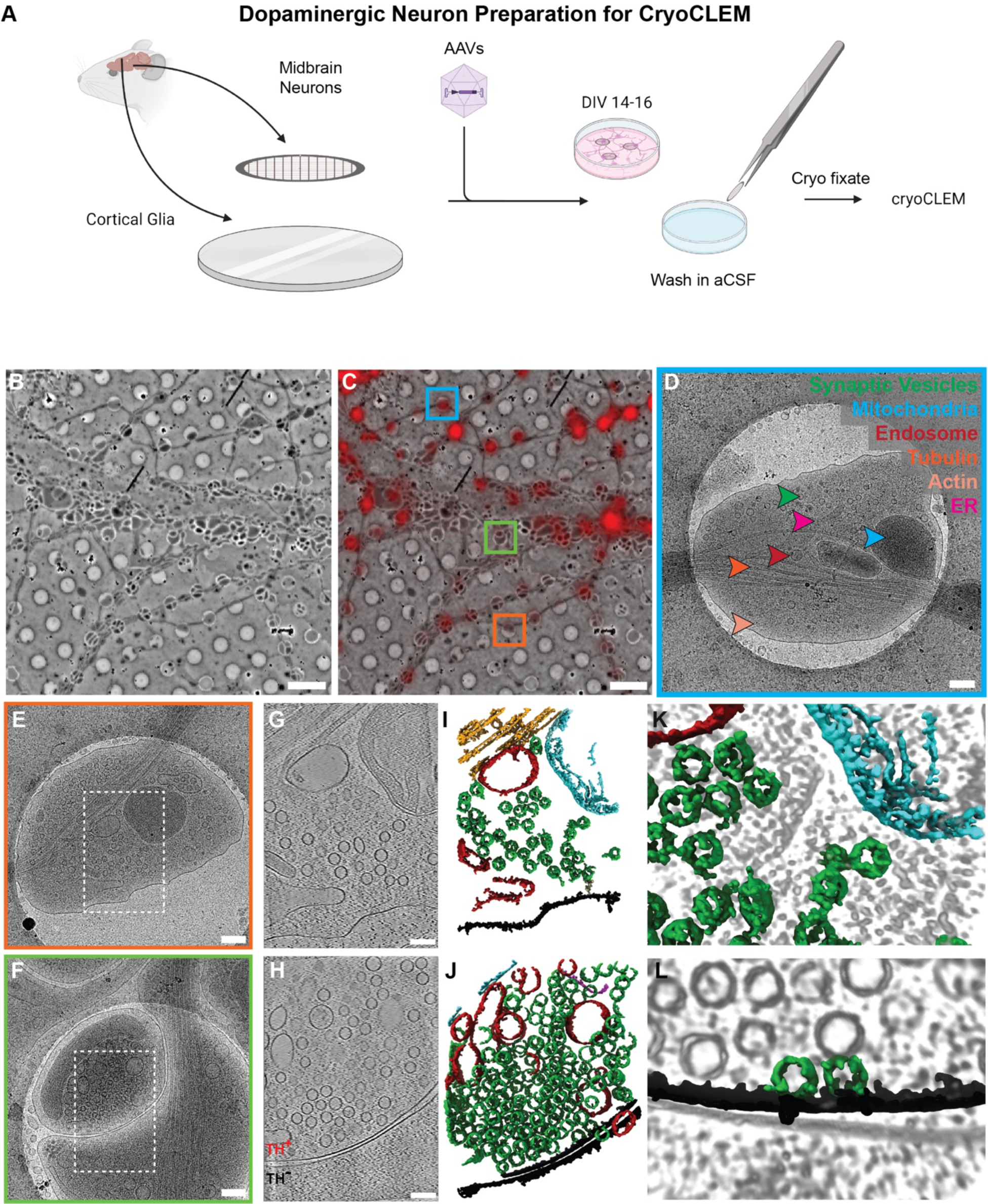
CryoCLEM imaging of DA presynaptic release sites using the Cell-Grid-Cell preparation. **A)** Preparation of primary DA neuronal co-cultures, in which the cryoEM grid is sandwiched between a support layer of cortical glia cells and midbrain neurons. The culture is treated with two AAVs resulting in fluorescence signal originating from TH positive DA presynaptic release sites. The cryoEM grids are treated in artificial cerebrospinal fluid with the potential of adding DA neuron specific pharmacology prior to cryo fixation. **B-C)** CryoEM 2D Tile scans without and with fluorescence signal overlay for identifying DA neurons. Scale bars 5 µm. **D)** View of a single DA varicosity identified from 2D tile scans. Organelles depicted by arrows: Synaptic vesicles (green), mitochondria (cyan), endosomes or lysosome (red), microtubuli (orange), ER (magenta) and actin (salmon). Scale bar 200 nm. **E)** Isolated DA presynaptic release site **F)** Contacting DA presynaptic release site. Scale bars 200 nm **G, H)** Slices through tomograms from images E and F (white insert). TH-positive (TH^+^) and TH-positive (TH^−^) varicosities indicated in **H**. Scale bar 100 nm **I, J)** Segmentations of the tomograms in G and H. Color scheme as above **K)** Close-up of synaptic vesicles and a mitochondrion in the cytosol. **L)** Close-up of two presynaptic vesicles in contact with the plasma membrane.

### The ultrastructure of DA presynaptic release sites

To describe the ultrastructure and composition of DA presynaptic release sites we collected both medium resolution 2D tile scans (pixel size of 12.5 Å) and high-resolution tilt series for 3D tomographic reconstruction (Figure 1D-L). The appearance of the DA neuronal extensions ranges from highly bundled, branching structures to individual extensions with varicosities displaying variable levels of TH expression (Figure S3A-B). In general, we see preference for formation of varicosities within the 2 μm spaced 2 μm diameter CryoEM grid holes, which has also been observed in the absence of a supporting cell layer for hippocampal and DRG neurons^18,19^. A previous study on nucleus accumbens DA presynaptic release sites in slices found a mean varicosity diameter of 0.4 μm and inter-varicosity distance of 1-5 μm^11^, suggesting that the dimensionality of the cryoEM grid (being an artificial substrate) is not necessarily restricting the frequency and size of the DA presynaptic release sites observed here.

At the organelle level, we detect the presence of microtubule bundles (1-8 microtubuli) travelling through the varicosities like molecular highways, branching actin cytoskeleton, mitochondria, endoplasmic reticulum (ER), endosomal compartments, presynaptic vesicles and rare events of molecular vaults (Figure 1D-F, S4). The presynaptic vesicles are relatively homogenous in size (46 +/− 6 nm, n=780) as gauged from the tomograms and are clearly distinguishable from the more tubular ER and larger and irregularly shaped endosomal compartments (Figure 1G-J). In our cultured neurons we find examples of both isolated and synapse-like contacting DA presynaptic release sites (Figure 1D-H, S3C-D). While the organelle content is similar, the distribution within synapse-like contacting DA presynaptic release sites appear more polarized with the presynaptic vesicles located close to the plasma membrane contact site and the mitochondria and microtubuli more distally located. In cases were the DA presynaptic release sites (TH-positive) and TH-negative extensions can be clearly discriminated, we find the TH-negative varicosities to be generally less crowded and less enriched in mitochondria and vesicles. Throughout the cultures we do not see indications of electron dense postsynaptic regions within the TH-negative varicosities when in contact with DA presynaptic release sites, but we do observe plasma membrane invaginations and vesicle fusions within the DA presynaptic release sites (Figure 1J,L, S5A).

The appearance and distribution of mitochondria is one of the most prevalent features. All DA presynaptic release sites identified in our cultures contains at least one mitochondrion which vary greatly in size (mean area: 3682 nm^2^, max area = 24429 nm^2^, n=295) and shape (mean circularity: 0.57, range: 0.1 – 0.91, n=295) (Figure 2A-E). We occasionally see exvaginations of the mitochondrial outer membrane that could potentially be intermediates in mitochondria-derived vesicle (MDV) formation (Figure 2F-G) ^20^. MDVs have been identified to operate under baseline conditions but also as a first line defense against stress in cardiac cells^21^. Their occurrence on synaptic mitochondria could therefore be a mechanism to reduce the negative impacts from their high oxidative phosphorylation activity. In the inner lumen of the mitochondria, we observe varying levels of calcium phosphate granules which have been suggested to serve as a repository for mitochondrial calcium and phosphate and are known to be dynamic in neuronal mitochondria (mean granule diameter: 22.84 nm, stdev: 14.46 nm, mean granule fractional occupancy of mitochondria area: 9.00%, stdev: 8.83%)^22^.

**Figure 2.**
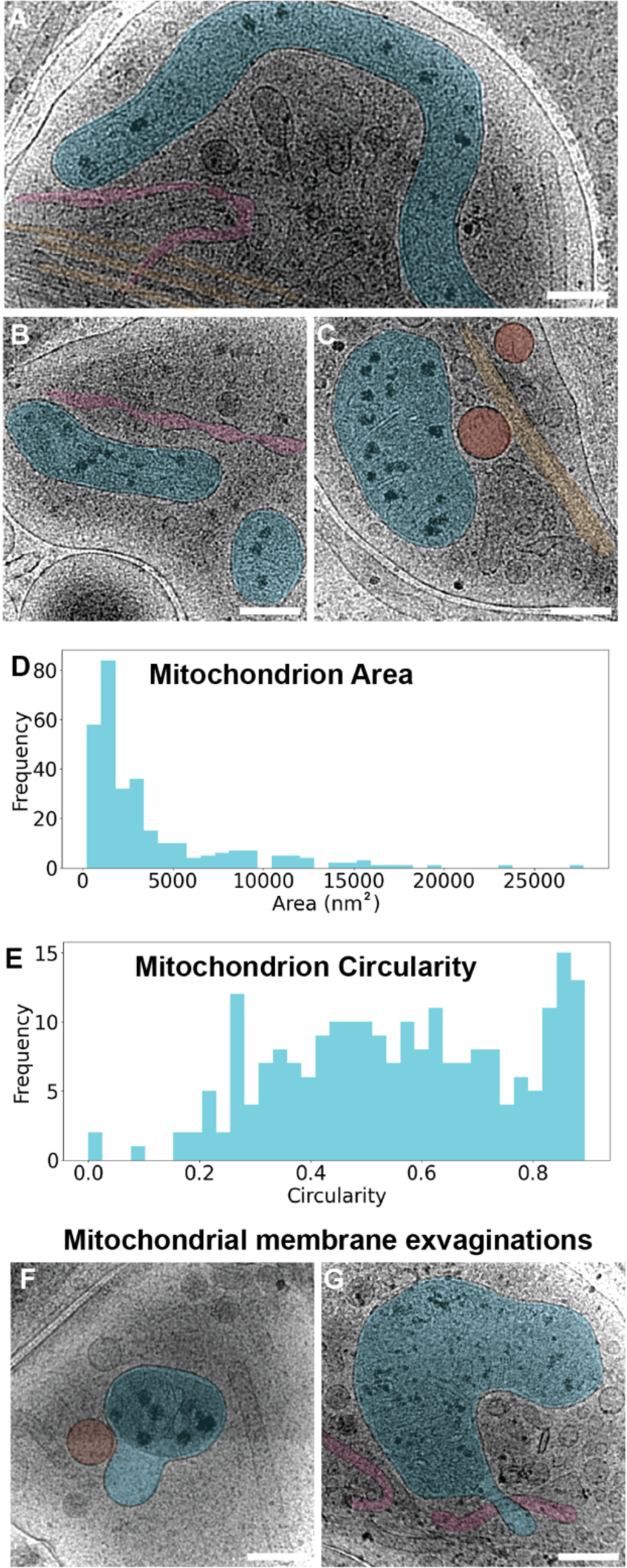
Mitochondrial variation within DA presynaptic release sites. **A-C)** Representative. examples of variation in mitochondrial (cyan) size, shape, and calcium phosphate deposits within DA presynaptic release sites. Quantification of total area **(D)** and circularity **(E)** of individual mitochondria. **F-G)** Examples of mitochondrial membrane exvaginations. ER and endosomal compartments depicted in magenta and red, respectively. Scale bars 200 nm.

### CryoET allows for single protein detection and in situ reconstructions of TRiC/CCT and the V-ATPase

The reconstructed tomograms allow for direct visualization of densities originating from larger proteins and protein complexes (typically >500 kDa). These include inter-organelle contacts linking the mitochondria to the ER, endosomes and microtubule^23,24^ (Figure 3A-C) and repetitive protein structures such as the microtubule inner proteins (MIPs) and actin^15^ (Figure S5B-D). Interestingly, we do not in our tomograms detect ribosomes, which have been observed in both cortical and hippocampal neuronal presynaptic sites imaged at similar resolutions^25,26^. In the plasma membrane we observe electron dense areas which may resemble smaller transmembrane (TM) proteins such as the 12 TM DA transporter (DAT), the 7 TM DA Receptor (D2R), or acetylcholine receptors (AchR). While these are difficult to visually separate, voltage gated K^+^/Na^+^/Ca^2+^ channels have a distinctive 25 TM tetrameric shape of ∼100-120 Å in diameter that can be observed in the tomograms (Figure 3D). In the presynaptic cytoplasm we also recognize the TRiC/CCT as round shapes with a diameter of ∼150 Å (Figure 3E). TRiC/CCT is an ATP dependent chaperone assisting in the folding of ∼10% of the proteome with actin and tubulin being some of its best-known substrates^27,28^. Using template matching and subtomogram averaging we detected a total of 459 TRiC/CCT particles in 59 tomograms from which we resolved a low resolution *in situ* structure. The structure resembles the closed TRiC/CCT configuration consisting of 16 CCT molecules arranged into two half spheres related by two-fold symmetry (Figure 3F). We do not see any apparent clustering or organelle association of the TRiC/CCT particles (Figure 3G). We also recognized larger vesicle membrane associated structures with the size and shape resembling the V-ATPase (Figure 3H). The V-ATPase is responsible for establishing the pH gradient across the membranes of synaptic vesicles and endosomal compartments. In DA neurons, the V-ATPase indirectly drives the proton dependent transport of neurotransmitters over the vesicular membrane via the vesicular monoamine transporter 2 (VMAT2)^29^. We averaged and aligned the vesicle membranes and created a centered elliptical shape as a decoy for subsequent template matching and subtomogram averaging. This approach identified 519 particles which could be refined into a low resolution *in situ* structure of the V-ATPase (Figure 3I). From the density resolved here we can distinguish the head (subunits A and B), the central stalk (subunits D and F), and the collar (subunits H, a-NTD and C) of the V_1_ domain. We see 1-2 V-ATPase molecules per vesicle within the DA presynaptic release sites (Figure 3J).

**Figure 3.**
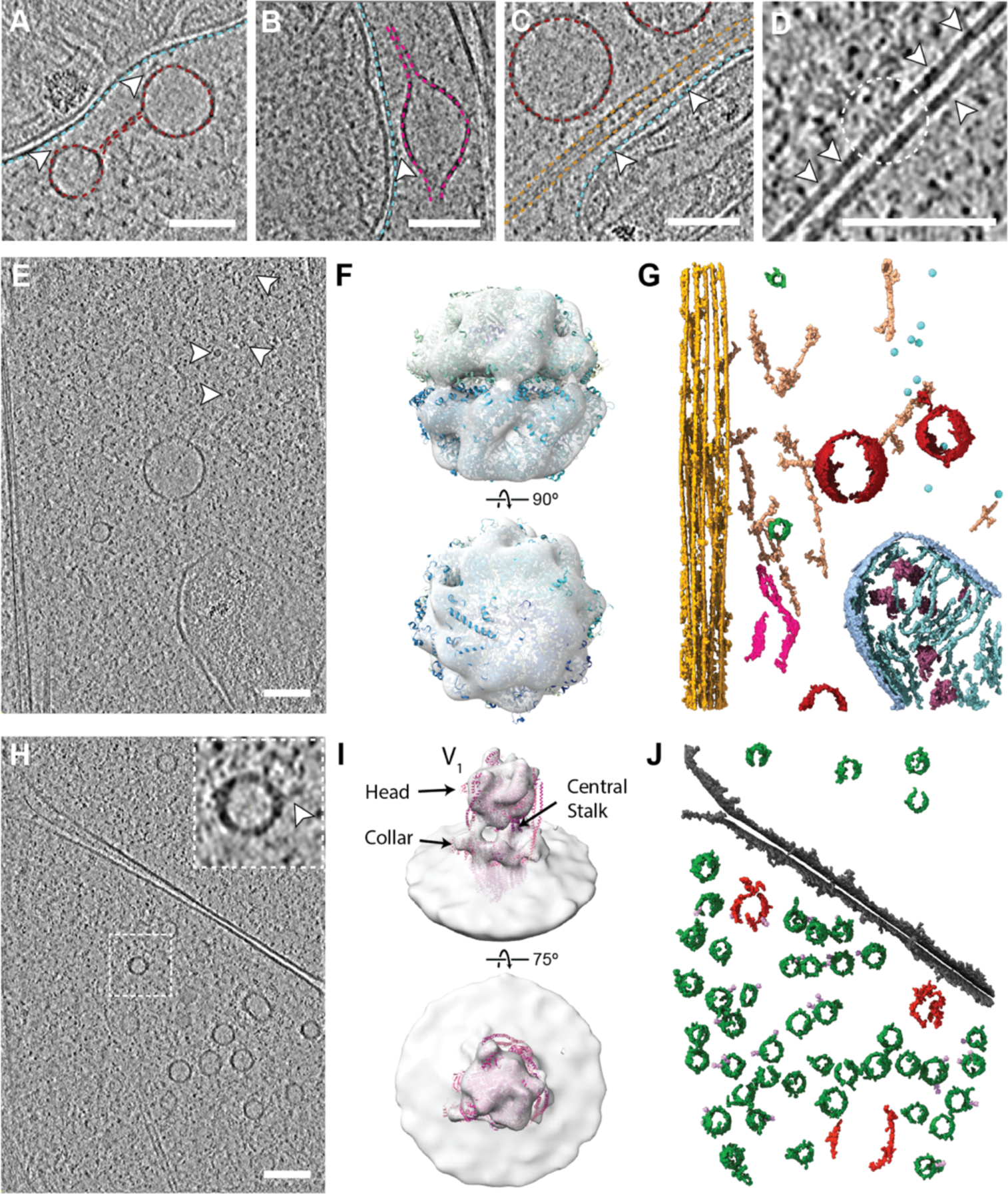
CryoET identifies inter-organelle contacts and allow for subtomogram averaging to resolve *in situ* structures of TRiC/CCT and the V-ATPase. **A)** Slice through tomogram showing contacts between mitochondrion and endosomal compartments. The mitochondrial outer membrane is depicted by cyan dashed line and the endosomal compartments by red dashed lines. White arrows indicate protein density with contact to both compartments. **B)** Slice through tomogram showing contacts between mitocondrion and ER (dashed line in magenta). White arrow indicates protein density with contact to both compartments. **C)** Slice through tomogram showing contacts between mitochondrion and microtubule. The microtubule edges are depicted by orange dashed lines. White arrows indicate protein density with sites of contact. **D)** Four-fold axis side view of a voltage gated K^+^/Na^+^/Ca^2+^ channel. The structure measures ∼100-120 Å in diameter. White arrows indicate density originating from other membrane proteins that are too small to identify **E)** Slice through tomogram with white arrows depicting TRiC/CCT chaperones measuring ∼150 Å in diameter. **F)** Reconstructed TRiC/CCT density obtained *in situ* with high resolution structure (PDB ID: 7LUP) fitted into the obtained density (see also figure S6). **G)** TRiC/CCT particles inserted back into segmented tomogram. Detected TRiC/CCT particles seems to be slightly clustered but primarily cytosolic and not anchored. **H)** Slice through tomogram with white arrow depicting V-ATPase stalk and head density on the surface of a presynaptic vesicle. **I)** Reconstructed V-ATPase density obtained *in situ* with high resolution structure (PDB ID: 6XBW) fitted into the density (see also figure S7). **J)** V-ATPase density inserted back into segmented tomogram. We see maximally two V-ATPase molecules per synaptic vesicle. Scale bars 100 nm.

### The ultrastructure of DA presynaptic release sites dynamically changes in response to pharmacological treatment

Finally, we exposed our primary DA cultures to pharmacological treatments and assessed the effect on the ultrastructure of the DA presynaptic release sites. The neuronal cultures were exposed for two minutes to artificial cerebral spinal fluid (aCSF) containing either 100 nM haloperidol or 10 µM DA (Figure 1A). There is tonic DA release by these neurons in this preparation and the neurons have presynaptic inhibitory DA D2 atuoreceptors^7^. Activation by DA of these D2 autoreceptors reduces firing of DA neurons as part of a negative feedback loop^30^. Haloperidol, a DA D2 receptor antagonist, will therefore promote firing of the DA neurons by impeding the negative feedback loop^7^. Through this pharmacological paradigm we were able to examine increases and decreases in firing at the DA presynaptic release sites. We obtained 2D tile scans of treated DA neurons and hand-segmented synaptic vesicles and mitochondria within the DA presynaptic release sites (Figure 4A-C). Vesicle density was assessed by comparing the number of vesicles within a 1 µm radius of each vesicle through kd-tree based neighbor identification. Mitochondria size was determined by area in 2D. From these quantifications, we observed that under haloperidol treatment conditions, and thus presumably during higher neuronal firing activity, there is an increase in the presynaptic vesicle density while DA treatment reduces the vesicle density (Figure 4H). As for mitochondrial size, there is an increase following DA treatment but no significant effect from haloperidol (Figure 4I). We also assessed the levels of calcium phosphate granules as a function of both treatments. Granules were identified through a random forest classifier, which allowed for determination of area fraction and diameters of calcium phosphate granules within a given mitochondrion (Figure 4D-G). DA treatment resulted in a modest decrease in the area fraction of granules and no change in the granule diameter, while haloperidol resulted in a dramatic reduction of the area fraction of granules as well as the granule diameter (Figure 4J-K).

**Figure 4.**
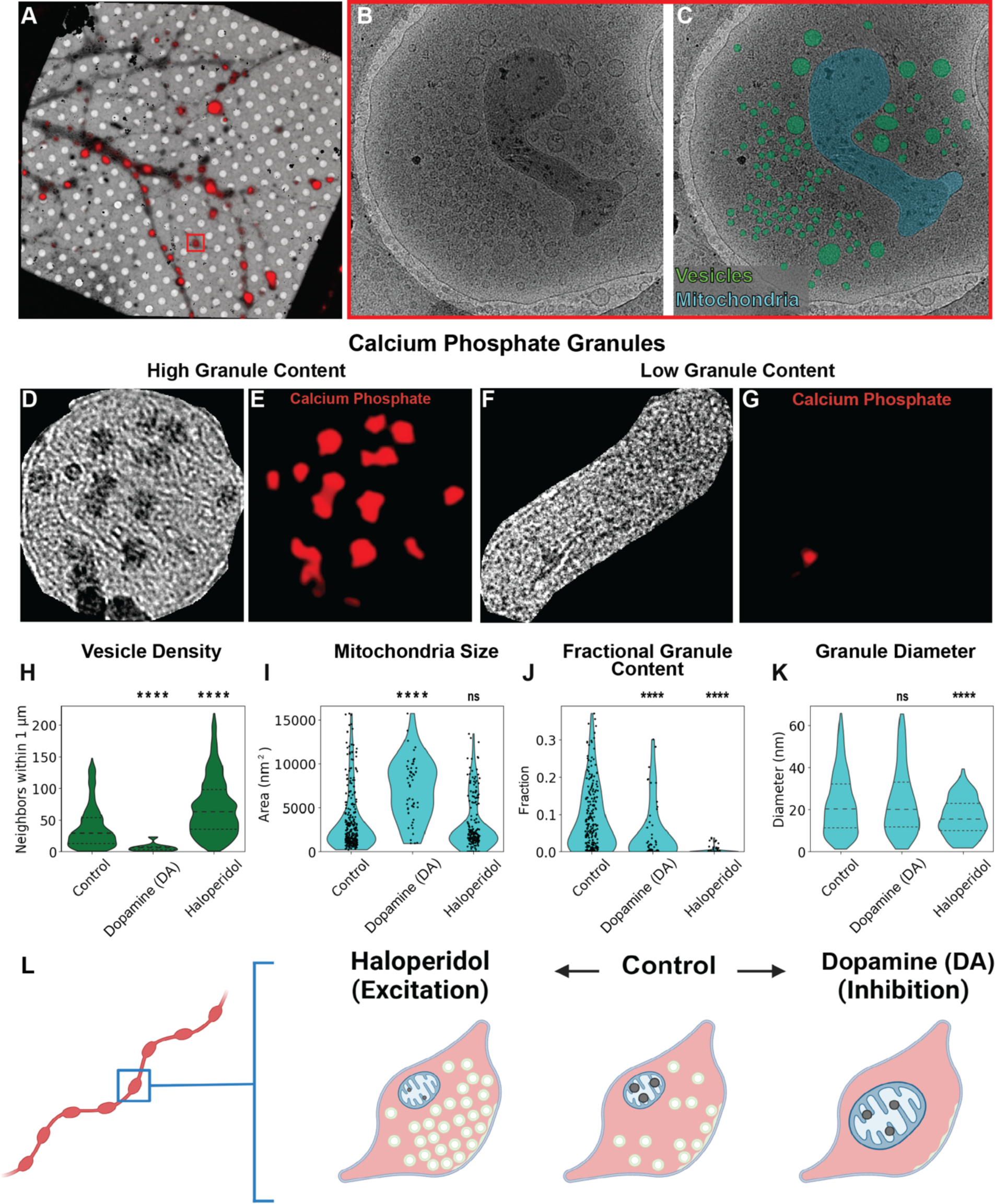
Exposure to DA or haloperidol bidirectionally modulates vesicular content and influences mitochondrial size and calcium phosphate deposition. **A)** 2D tile scan of cryoEM grid square containing TH-positive DA presynaptic release sites. **B-C)** Representative close-up of a DA presynaptic release site showing an example segmentation for vesicles (green) and mitochondrion (cyan). Similar segmentations were performed within samples threated with either control aCSF, DA (10 µM), or haloperidol (100 nM). **D-G)** Calcium phosphate granules within mitochondria were identified with a trained random forest classifier. Example segmentations are shown for two mitochondria, one with many granules and one with just a single granule. The pixel classifier was able to distinguish artifacts from the calcium phosphate granules (bottom of image D). **H)** The number of neighboring vesicles within 1 µm from each vesicle for each of the pharmacological treatments. Number of segmented vesicles in the datasets are: 4258(control), 256(DA) and 6851(haloperidol). DA p value < 0.0001, Haloperidol p value < 0.0001. **I)** The area of mitochondrion in nm^2^ as measured from the 2D tile scans for each of the pharmacological treatments. Segmented mitochondria in datasets are: 295 (control), 51(DA), and 199(haloperidol). DA p value < 0.0001, Haloperidol p value > 0.1. **J)** The ratio of pixels that were identified as being calcium phosphate granules for each mitochondrion per pharmacological condition. DA p value < 0.0001, haloperidol p value < 0.0001. Number of zero value entries per condition: 9 (control), 7 (DA), 92 (haloperidol). **K)** Comparison of the diameter of each granule. DA p value >0.5, Haloperidol p value < 0.0001. **L)** Summary of the ultrastructural changes within DA presynaptic release sites as a function of either DA or haloperidol.

## Discussion

The Cell-Grid-Cell method developed here relies on separating a stimulatory and/or inhibitory supporting cell layer from the cells of interest which are seeded on top of the cryoEM grid for subsequent cryo fixation. The methodology could be generalized to allow for imaging of a large range of complex cellular systems where isolation of one or more cell types is desirable e.g. immune cell contacts, targeted viral infections or host-parasite interactions.

In this study, we used the Cell-Grid-Cell method to separate a supporting cortical glial cell layer stimulating development and axonal growth of midbrain neurons. DA neurons were identified via positive fluorescent labelling. In our cultures, we find that the DA neuronal extensions adopt different overall morphologies ranging from highly bundled to individual axonal extensions but with well-defined TH-positive DA presynaptic release sites. In the cultures, we observe many interfaces between DA presynaptic release sites and TH-negative varicosities. The TH-negative varicosities display structures and organelle content more similar to postsynaptic structures observed in hippocampal glutamatergic neurons in cultures^13,31^, however, we do not observe postsynaptic densities (PSDs), which have otherwise been described for both hippocampal glutamatergic and GABAergic neurons imaged at similar resolutions^14,31^. This finding is in good agreement with a recent EM study of DA neurons in sections of mouse nucleus accumbens, which found a majority of DA axonal varicosities to show no evidence of widespread structural connections with other neurons, and only rarely engage in classical synaptic transmission with clear PSD in the target neuron^11^. These observations are furthermore supported by previous studies indicating that DA synapses are not described by classical synaptic models but more often engage in volumetric neurotransmission^32–35^.

Within the DA presynaptic release sites, we focused our investigations on presynaptic vesicles and mitochondria. In the untreated DA neurons, the presynaptic vesicles are relatively homogenous in size but vary in numbers. The mitochondria are spatially prevalent but vary greatly in appearance and in the content of calcium phosphate granules. DA neurons are typically unmyelinated and have been shown to extensively arborize and chronically fire with pacemaker activity which result in an increased need for synaptic mitochondria, thus making DA neurons particularly impacted by mitochondrial targeting stimuli^36,37^. To test the ultrastructural dynamics of these two organelles we exposed the cultures to brief pharmacological treatments prior to cryo-fixation. DA, an agonist on presynaptic DA D2 receptors, causes inactivation of dopaminergic neurons, while haloperidol, a DA D2 receptor antagonist, causes activation of DA neurons by blocking binding of tonically released DA in the culture (Figure 4L)^7,30^. For the presynaptic vesicles, we observed an increase in numbers in the activated state and a decrease in the inactivated state compared to the untreated control. This suggests that a reserve pool of presynaptic vesicles is not necessarily maintained but also that the vesicle formation process occurs at a faster rate than the vesicle release process. The delay in vesicular release could potentially play an important role in regulating the vesicular DA content. The mitochondria were found to be larger in the inactivated neurons but unchanged in the activated neurons. In both cases, the fractional area of the granules decreases. While the increase in mitochondrial size explains the decrease in the granule fractional area in the inactivate state, the activated neurons display a dramatic reduction of the granule fractional area and diameter which suggests that mitochondrial calcium phosphate is indeed involved in regulation of DA neuronal firing. The rapid reshaping of the mitochondria in our cultures is in good agreement with previous findings showing that mitochondrial structural changes alter mitochondrial functional capacity^38,39^, and that DA neurons in general have been found to be particularly vulnerable to impaired mitochondrial function induced by pharmacology such as MPTP, rotenone or cocaine, or genetically through alterations of PINK1 or Parkin^11, 40–43^.

At the single protein level, we used the high-resolution tomograms to identify protein complexes involved in mitochondrial contacts with endosomes, lysosomes, and ER. Mitochondria-endosome contacts are interesting in the context of potential iron transfer which is necessary for oxidative phosphorylation^44,45^. *In vivo,* depletion of iron uptake into DA neurons yielded damaged mitochondria, axonal injury and phenotypical PD symptoms^46^. Mitochondria-lysosome contacts form dynamically for the exchange of molecular components and are found to influence the mitochondrial degradation process^47^. In PD patients, impaired mitochondria-lysosome contacts have been found to severely disrupt mitochondrial function^48^. ER-mitochondria contacts have been suggested to facilitate transfer of calcium from ER to mitochondria^49,50^. It was previously found that reduction of ER-mitochondria contacts in DA neurons, through an α-synuclein dependent mechanism, disrupts mitochondrial ATP production, demonstrating the vital role of this inter-organelle contact in the development of PD^51^.

From our tomograms we were also able to resolve low resolution *in situ* structures of TRiC/CCT and the V-ATPase. TRiC/CCT is a key chaperone shown to bind and promote folding of ∼10% of the eucaryotic proteome^52^. Through cell type specific subcellular proteomics, it has been shown that TriC/CCT is axonally localized in DA neurons and maintains correct synaptic protein structure^3^. Protein misfolding in DA neurons is a hallmark of DA neuron specific neurodegenerative disorders (i.e PD), where aggregation and fibrillation of α-synuclein (Lewy bodies) are often detected in DA neurons, postmortem. Fibrillation of α-synuclein has been shown to be directly inhibited through the function of TRiC/CCT^53^, and therefore further studying the distribution of this complex under pathological conditions could be of significant interest. The V-ATPase is vital to DA neuronal function and facilitates DA vesicular uptake through the vesicular monoamine transporter 2 (VMAT2) that is powered by a proton gradient across the membrane of synaptic vesicles^29^. This gradient is maintained through the proton pumping activity of the V-ATPase, creating an internal vesicular pH of ∼5.6^54^ which is sufficient to maintain approximately 33,000 DA molecules within each synaptic vesicle^55^. In glutamatergic hippocampal neurons, the V-ATPase has been estimated to be present in numbers of 0.7 to 2 copies per vesicle^56^ which is in good agreement with our findings of maximally 2 V-ATPase molecules per presynaptic vesicle. Psychostimulant drugs, such as amphetamine, operate in DA neurons through the alteration of synaptic vesicle pH and therefore understanding the distribution of the V-ATPase under such conditions could potentially aid in further understanding their mode of action^57^.

In summary, this study offers a methodology for description and perturbation of the ultrastructure and dynamics of DA neurons. A general understanding of the ultrastructural dynamics within cultured DA neurons serves as an important platform for understanding neurophysiological disorders and the effect of pharmacological intervention in a more complex context.

## Materials and Methods

### Cortical Glia Preparation

Cortical glia cells were isolated from P1-P3 Wistar rat pups (Charles River, Germany)^7^. The brains were isolated from each pup and held in phosphate-buffered saline (PBS) on ice. Each cortex was cut from the brain, followed by the removal of the hippocampus and the meninges. The remaining tissue was cut into minimally sized sections and stored in dissection medium (Hanks’ Balanced Salt Solution with added Sodium Pyruvate (1 mM), Penicillin-Streptomycin (Sigma P0781, 200 U/L), 10 mM HEPES, and 0.54% glucose) on ice. Following the collection of all cortical sections, the dissection medium was aspirated away and replaced with glia cell media (DMEM with HEPES “1965”, FBS (10%), Penicillin-Streptomycin (60 U/L)). To form a single cell suspension, the tissue was titrated through flame-polished Pasteur pipettes. The cell suspension was then filtered through a 70 µm cell strainer and then underwent centrifugation for 10 minutes at 200-300 x g. The supernatant was removed and the pellet was suspended, filtered through a 70 µm cell strainer, and centrifuged a second time at 200-300 x g. The pellet, containing the cortical glia cells, was suspended in cortical glia cell media and grown in large T175 flasks, where 1.5 to 2 brains were used per flask. The cortical glia cells were grown in an incubator at 37° C with 5% CO2. Upon achieving 70% confluency, the cells from a single large T175 flask were then split across two large T175 flasks, and grown until 70% confluency. Cells were removed from the flask through trypsinization, and isolated through centrifugation for 10 minutes at 200-300 x g. The cells were resuspended in heat-inactivated serum with 10% DMSO, and the cells from a single large T175 flask was divided into 4 separate aliquots and placed in a Corning® CoolCell® LX container at −80° C. Following completion of freezing the cells were stored under liquid nitrogen until use.

Two weeks prior to a given dissection, an aliquot of frozen glia cells was thawed, and grown in glia media at 37° C with 5% CO2 for one week in a large T175 flask. One week prior to the dissection glia (50,000 cells/mL) were added to each well of a 6 well tray. Each well in the tray, prior to the addition of cells, was washed first with ethanol (95%), and incubated with poly-D-lysine (10 mg/mL) for 2 hours at 37° C. Wells were then washed 5x with H_2_O to remove unbound poly-D-lysine, and glia cells were added. On the day prior to the addition of neurons, the media was replaced with DA neuron media (Neurobasal A with GlutaMAX (1/100), B-27+ (1/50), ascorbic acid (200 μM), Penicillin-Streptomycin (60 U/L), and kynurenic acid (0.5 mM).

### Dopamine (DA) Neuron Production

Midbrain DA neurons were isolated from P1-P3 Wistar rat pups (Charles River, Germany)^7^. The brains were first isolated from each pup and stored in dissection medium (Hanks’ Balanced Salt Solution with added Sodium Pyruvate (1 mM), Penicillin-Streptomycin (Sigma P0781, 200 U/L), 10 mM HEPES, and 0.54% glucose) on ice. The midbrain of each brain was then cut by scalpel under the view of a dissecting microscope, in order to take the section likely containing the ventral tegmental area and the substantia nigra pars compacta based off anatomical landmarks. These sections were cut into millimeter sized pieces and placed in papain solution (cysteine (1 mM), DNase (0.1 mg/mL), papain (20 units/mL), NaCl (116 mM), KCl (5.4 mM), CaCl_2_ (1.9 mM), NaHCO_3_ (26 mM), NaH_2_PO_4_H_2_O (2 mM), MgSO4 (1 mM), EDTA (0.5 mM), Glucose (25 mM), kynurenic acid (0.5 mM) in water) for 30 min at 37◦C and 5% CO_2_ 95% O_2_. The media containing the tissue was then replaced with pre-warmed DA neuron media. The tissue was titrated with flame-polished Pasteur pipettes, and the suspension was centrifuged at 200-300g for 10 minutes. The pellet was suspended in DA neuron media, and cells were added at 50,000 cells/mL to each well. Glia-derived neurotrophic factor (GDNF) (100ng/mL) was added to the cultures. Three days later 5-Fluorodeoxyuridine (FDU)-solution (Uridine 16.5 mg/mL, 5-FDU 6.7 mg/mL) was added to each culture at a 1:1000 dilution to inhibit glia cell proliferation.

### Preparation of Cryo-EM Grids

Quantifoil® R 2/2 Au 200 grids were glow discharged with a Leica Coater ACE 200 for 15 mV for 30 seconds. Next, they were washed once with PBS, and then incubated in with poly-D-lysine (20 mg/mL) for 2 hours at 37° C. Grids were then washed 5x with PBS, and stored in PBS until use. Grids were placed on the layer of glia cells immediately prior to the addition of DA neurons.

### Viral Transduction

Viral production was carried out as previously described^7^. An in-house constructed pAAV-pTH-iCre-WPREpA vector was developed, featuring a truncated 488bp rat TH promoter that encodes the Cre recombinase. Recombinant AAV2retro-hSyn-FLEX tdTomato viruses were also generated in-house, employing the FuGene6 (Promega) facilitated triple plasmid co-transfection technique in HEK293t cells. Post three days of transfection, the cells underwent harvesting, and the virus was purified utilizing a modified Iodixanol gradient purification method. The genomic titer of the AAV was ascertained using a PicoGreen-based technique. The expression of Cre-dependent tdTomato mediated by AAV2-retro-pTH-iCre was authenticated through fluorescence microscopy, revealing no discernible differences across the various titers utilized. For the process of transduction, the virus was introduced to primary culture DA neurons at a rough concentration of 2*10^9^ vg/mL, between 1 to 3 days following the preparation of neuronal cultures.

### Pharmacological Treatment

Immediately prior to vitrification, cultures were first washed in artificial cerebral spinal fluid (aCSF) (NaCl (120 mM), KCl (5 mM), CaCl_2_ (2 mM), MgCl_2_ (2mM), HEPES (25 mM), glucose (30 mM), pH 7.4). To treat with pharmacology, samples were then placed in aCSF containing either 100 nM haloperidol, 10 µM DA, or control aCSF for 10 seconds prior to being placed in the vitrobot. Given the time between placing in the vitrobot and freezing, it was timed that the samples were exposed to the pharmacological treatment for 2 minutes

### Cryo Fluorescence Microscopy

Cryo-fluorescent imaging was conducted on a FEI CorrSight spinning disk confocal equipped with a cryostage (FEI). Detection was with a Hamamatsu ORCA-Flash 4.0 sCMOS camera with an EC Plan-Neofluar 40x/0.9 NA Air objective. Tile scans were collected with a z step of 200 nm to capture the entirety of the EM grid. Transmission imaging was collected as well as fluorescence excitation with 561 nm illumination. Images were stitched with the MIST imageJ plugin^58^.

### Cryo Electron Microscopy (CryoEM)

Grids were lifted carefully out of the trays and manually backside blotted at 4°C, 100% humidity and plunge-frozen in liquid ethane using a Vitrobot Mark IV (Thermo Fisher). Initial grid screening and high-resolution tile scan montage micrographs were acquired using a Titan Krios G2 equipped with a Falcon III camera operated at 300 keV in linear mode. For the high-resolution montages, we used a nominal magnification of 6500kx resulting in a pixel size of 12.5 Å, a defocus of 20 μm and 3 sec exposure (1 e/Å^2^/sec). The tile scan montages were acquired using MAPS 3 software (Thermo Fisher) allowing for direct import and alignment of the stitched fluorescence light microscopy atlases obtained prior to acquiring high resolution maps. Images were stitched with the Imaris Stitcher software. For regions in which the stitching with Imaris Stitcher was unsuccessful, BigStitcher was utilized ^59^. Image registration to fluorescence images was performed manually applying rotation, scaling, and translation.

### Analysis of 2D images

Varicosities were positively identified as dopaminergic through visualization of the correlated data from MAPS 2 software. Mitochondria and vesicles were segmented by hand through napari ^60^. Vesicle density and mitochondria size were measured through home written python scripts. Calcium phosphate granules were identified through a random forest classifier with ilastik^61^. The granule fraction and the diameter of the granules was computed through custom python scripts.

### Cryo Electron Tomography (CryoET)

CryoET tilt series were collected on a Titan Krios microscope (Thermo Fisher) operated at 300 kV using a BioQuantum energy filter (slit width 20 eV) and a K3 direct electron detector (Gatan). SerialEM was used for acquiring montaged maps for correlation with the cryo-FM maps and for tilt series acquisition^62^. Tilt series images were acquired at 2° angular increments over a ±60° interval in groups of 2 using a dose-symmetric tilt scheme at a nominal magnification of 53,000x resulting in an unbinned pixel size of 1.69 Å and a defocus range from −3 to −6.5 μm ^63^. Total exposure time was adjusted to maintain a total dose per tilt image of 2.4 e/Å^2^. Exposures were fractionated into 10 frames and saved as LZW-compressed TIFF images without normalization.

#### Processing

All movies were pre-processed using WARP 1.0.9 ^64^. Motion correction was performed with a temporal resolution of 10 for the global motion and 1 × 1 spatial resolution for the local motion. We considered motion in the 45.7–3 Å range weighted with a B-factor of −500 Å^2^. Micrographs displaying more than 5 Å intraframe motion were discarded. CTF estimation was performed using 8 × 5 patches in the 49-3.5 Å range. The raw unbinned tilt series were then exported for alignment in IMOD/etomo using patch tracking^65^. Briefly, the tilt series were coarsely aligned using low and high frequency rolloff sigma of 0.03 and 0.05 fractional reciprocal lattice units, respectively, and a high frequency cutoff radius of half Nyquist and with no Cosine stretching. The tilt series were binned by 4 before proceeding to fiducial model generation. The patch tracking was performed using 400×400 patches with a fractional overlap of 0.33. The resulting patch tracking alignment model was manually curated, and the fine alignment was performed by keeping tilt axis rotation, magnification, and tilt angles fixed. This alignment was then reimported back into WARP for estimating defocus and astigmatism for the entire tilt series and finally binned down to 13.52 Å/px (bin 8) and deconvoluted or denoised for inspection, segmentations and subtomogram averaging.

#### Subtomogram averaging

For TRiC/CCT, we used template matching as implemented in Warp^64^ and used a map prepared from PDB ID: 7LUP as reference ^66^. The map was low-pass filtered 40 Å before template matching. The matched particles were inspected using Cube function in Warp to select a Figure of Merit (FoM) cutoff of 10, which was then used to filter all particle lists before extracting 48^3^ subtomograms with a pixel size of 3.38 Å/px. The resulting particles were then imported into Relion 4 and subjected to several rounds of 3D classifications without alignment to filter out bad particles using the raw reconstruction from the particle list as template ^67,68^. To get rid of remaining particle artifacts we split the particle list to produce one file for each tomogram and manually curated all tomogram picks using ArtiaX as implemented in ChimeraX ^69^. We then matched the resulting files to the input files (to maintain particle file structure) and recombined them for importing back into Relion. This resulted in a final number of 459 particles in 52 tomograms (7 tomograms without any particles) which was refined using C2 symmetry and local searches of 0.9 degrees. The resulting particles and half maps were then imported into M for final refinement reaching a resolution of 25 Å. Processing steps are summarized in Figure S6.

For the V-ATPase, a first attempt to perform subtomogram averaging and classification on oversampled vesicle membranes did not yield good results, potentially due to low V-ATPase numbers. Instead, we prepared an average of the vesicle membrane surface using Dynamo^70^ and placed an elliptical decoy resembling the size of the V-ATPase in the center of the box which we then used for template matching template matching using in Warp. The matched particles were again inspected using Cube to select a FoM cutoff of 11, which was then used to filter all particle lists before extracting 56^3^ subtomograms with a pixel size of 6.76 Å/px. As for the TRiC/CCT subtomograms, we imported the particles into Relion and did several rounds of 3D classifications without alignment, as well as manual curation of the particles in ArtiaX. The resulting curated particles we subjected to 3D Refinement in Relion and was finally refined in M. Processing steps are summarized in Figure S7. The final reconstruction at a resolution of 34Å was compared to the V-ATPase structure with PDB ID: 6XBW^71^.

### Statistics

Statistics were completed by applying Mann-Whitney U tests between the experimental conditions and the control for each of the experiments.

## Acknowledgements

We thank Tillmann Hanns Pape, Pablo Hernandez-Varas, Cristiano Di Benedetto and Klaus Qvortrup for support during data collection at CFIM and Guillermo Montoya for access to the HPC infrastructure at the Novo Nordisk Center for Protein Research. We also thank Marta Carroni for access to the Krios Microscope at SciLifeLab via the CryoNET network. We thank Andreas Toft Sørensen and Søren Heide Jørgensen for the viral vector for dopaminergic neuron identification. This work was supported by the Novo Nordisk Foundation (S.E: NNF17OC0030788), the Lundbeck Foundation (MDL R230-2016-3154, UG R359-2020-2301,

UG R276-2018-792) and the Max Planck Society (J.A.G.B).

## Author Contribution

M.D.L and S.E., designed the project. M.D.L and S.E. prepared all samples and performed cryo-FM/EM/ET sample preparation/screening, analyses, reconstructions and subtomogram averaging. D.R.M assisted in acquisition of tilt series. M.D.L, S.E. J.A.G.B and UG discussed and interpreted the results. M.D.L and S.E, wrote the manuscript with input from D.R.M, U.G. and J.A.G.B.

