## Supplementary Information for "Ultrastructural Dynamics of Dopaminergic Presynaptic Release Sites revealed by Cryo-correlative Light and Electron Microscopy"

*Lycas et al.*

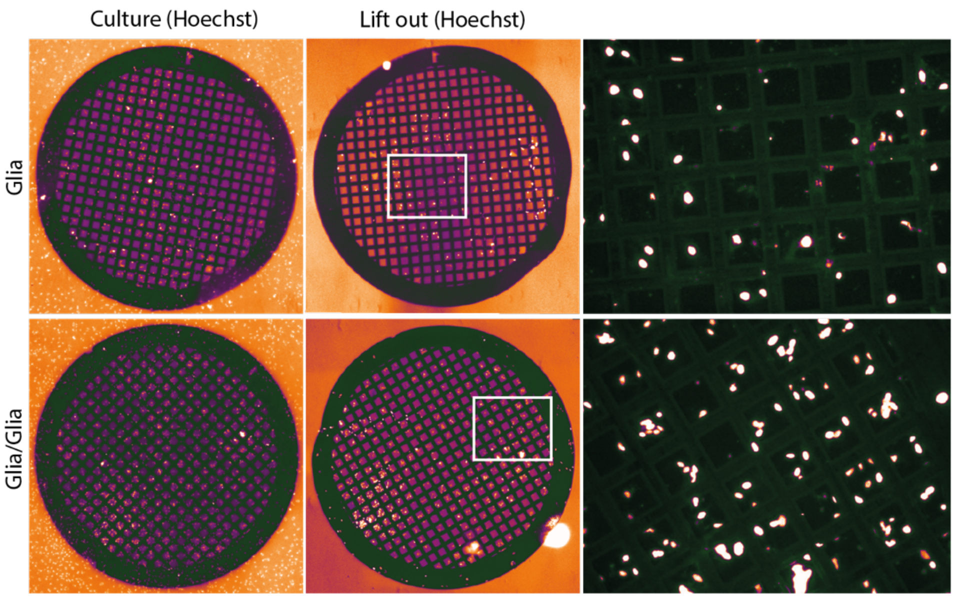


**Figure S1. Examination of the effectiveness of the Cell-Grid-Cell preparation.** The cryoEM grid was placed over the layer of a cortical glia support cell culture (top row). Layering of a second application of cortical glia cells on top (bottom row). Both sample preparations were allowed to grow for 14 days after which grids were lifted out of the culture dish and placed in an empty well. Cells were imaged by a Hoechst staining on a widefield fluorescent microscope.

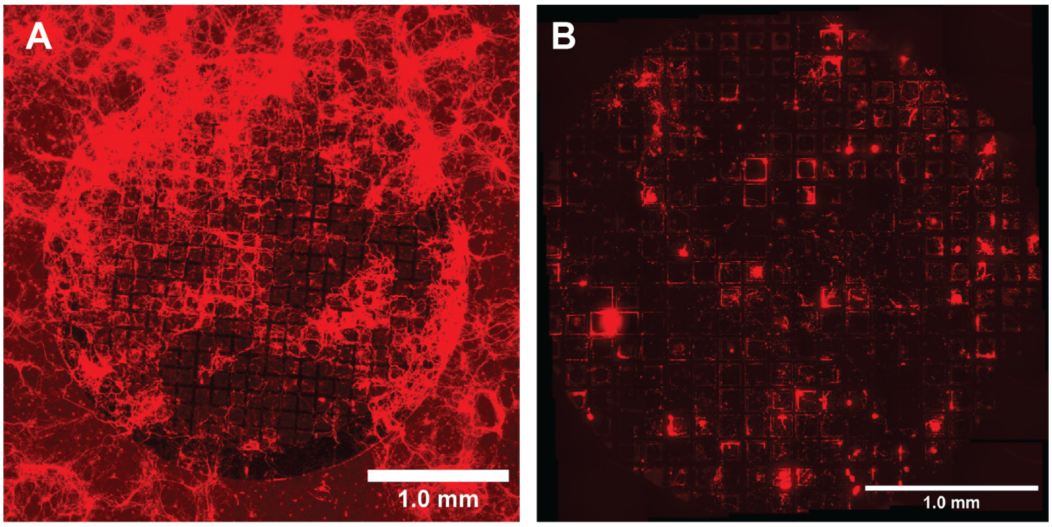


**Figure S2. Live and cryogenic imaging of DA neurons on cryoEM grid preparations.**
**A)** Live imaging by widefield fluorescence microscopy just prior to plunge freezing of cultured DA neurons expressing tdTomato. **B)** Cryo fluorescence microscopy image of the DA neurons on a (separate from image A) cryoEM grid following the plunge freezing procedure.

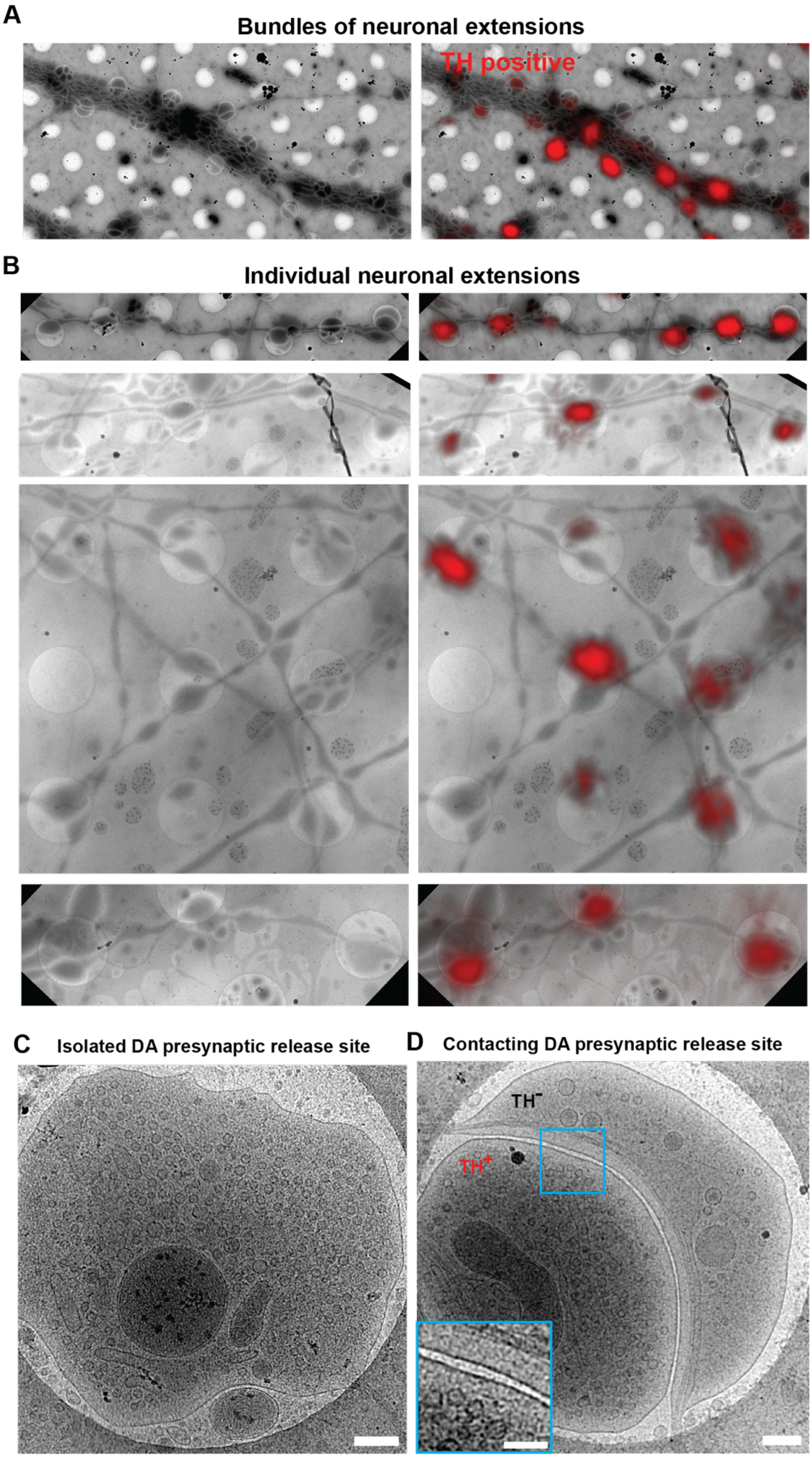


**Figure S3. The morphology and appearance of the TH positive and negative neuronal extensions A)** Highly bundled, branching TH-positive DA neuronal extensions intertwined with TH-negative extensions **B)** Individual neuronal extensions with TH-positive varicosities displaying variable fluorescence intensities and in context of different TH negative neuronal extensions and cellular cytosolic structures. **C)** Example of an isolated DA presynaptic release site. **D)** Example of a synapse-like contacting DA presynaptic release site. Close-up showing plasma membrane junction. TH-positive ($TH^+$) and TH-positive ($TH^-$) varicosities are indicated. Scale bars of C and D are 200 and 100 nm for close-up.

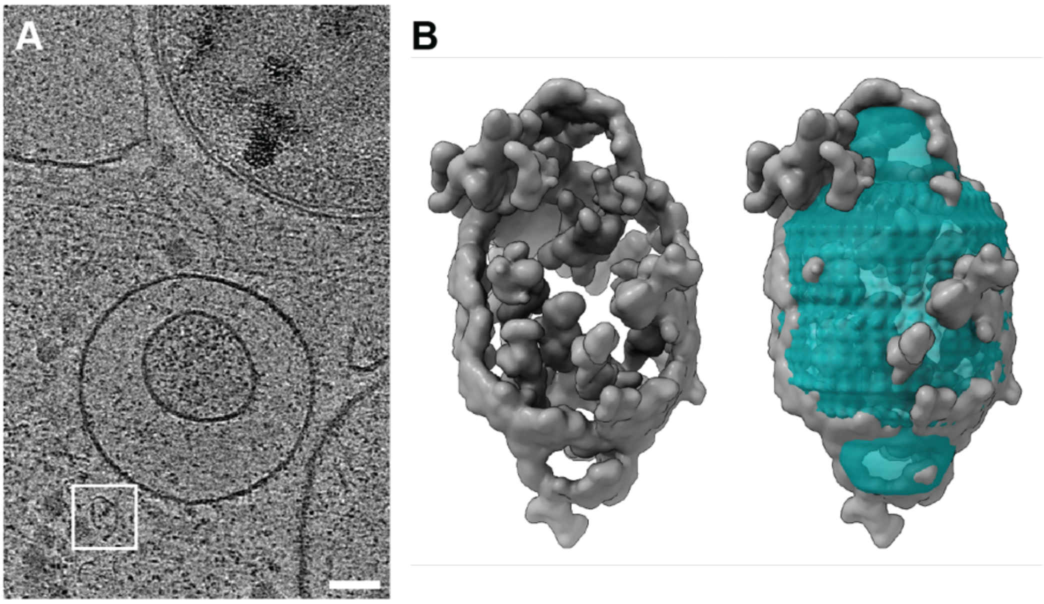


**Figure S4. Vault located in a DA presynaptic release site A)** Slice through tomogram with the identification of density corresponding to a Vault. Scale bar 50 nm. **B)** Extracted vault density from tomogram compared to the known structure of vaults (PDB ID: 4V60).

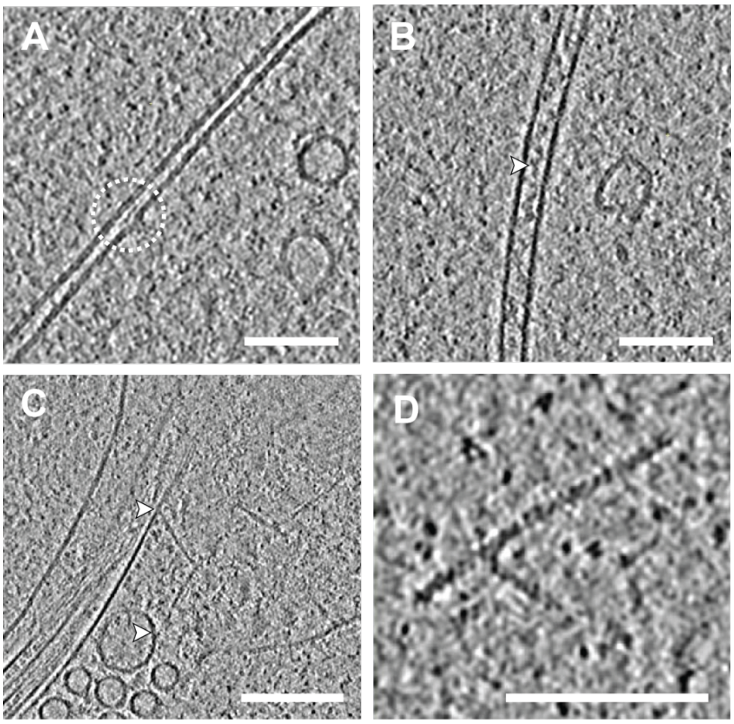


**Figure S5. Examples of membrane invagination and DA neuronal cytoskeleton. A)** Slice through a tomogram showing a membrane invagination. **B)** Slice through a tomogram showing a microtubule measuring ~25 nm in diameter, Microtubule inner proteins (MIPs) are clearly visible in the center (white arrow). **C)** Slice through a tomogram with an example of actin-plasma membrane and actin-organelle contacts (white arrows). **D)** Slice through a tomogram with an example of actin measuring ~60 Å in diameter with visible repetitive structure. Scale bars 100 nm

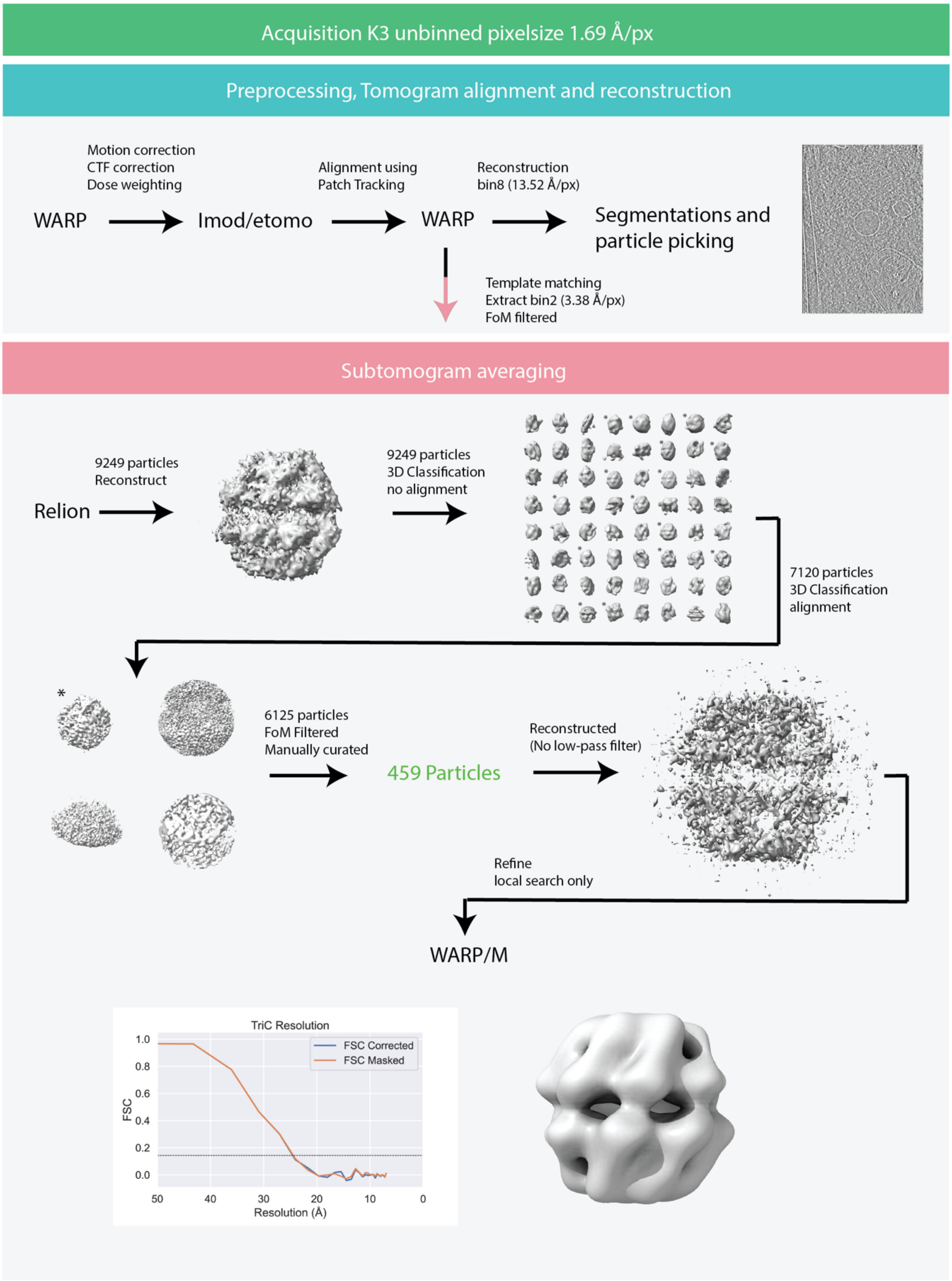


**Figure S6. Image processing flow charts for TRiC/CCT reconstructions.** For details see materials and methods. Asterisks indicate selected classes. Resolution of reconstructions are determined by gold-standard Fourier shell correlation (FSC) at the 0.143 criterion.

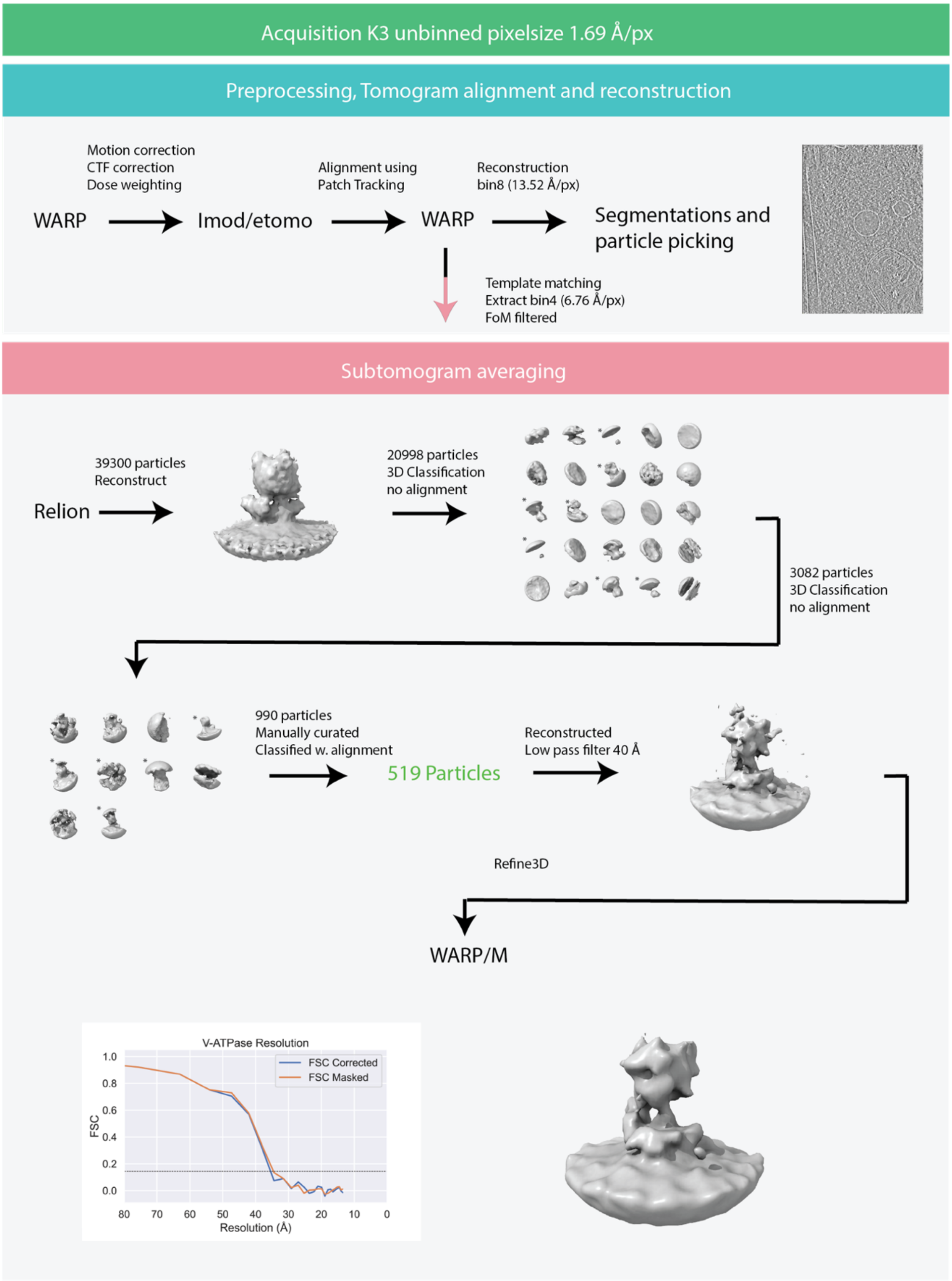


**Figure S7. Image processing flow charts for V-ATPase reconstructions.** For details see materials and methods. Asterisks indicate selected classes. Resolution of reconstructions are determined by gold-standard Fourier shell correlation (FSC) at the 0.143 criterion.
